# Palmitvaccenic acid (Δ11-cis-hexadecenoic acid) is synthesized by an OLE1-like desaturase in the arbuscular mycorrhiza fungus *Rhizophagus irregularis*

**DOI:** 10.1101/2020.01.13.901264

**Authors:** Mathias Brands, Edgar B. Cahoon, Peter Dörmann

**Author notes:** **Correspondence:** Peter Dörmann.

## Abstract

Arbuscular mycorrhiza (AM) fungi deliver mineral nutrients to the plant host in exchange for reduced carbon in the form of sugars and lipids. Colonization with AM fungi upregulates a specific host lipid synthesis pathway resulting in the production of fatty acids. The fungus *Rhizophagus irregularis* accumulates predominantly palmitic acid (16:0) and the unusual palmitvaccenic acid (16:1^Δ11cis^). Here, we present the isolation and characterization of *RiOLE1-LIKE*, the desaturase involved in palmitvaccenic acid synthesis, by heterologous expression in yeast and plants. Results are in line with the scenario that *RiOLE1-LIKE* encodes an acyl-CoA desaturase with substrate specificity for C15-C18 acyl groups, in particular C16. Phylogenetic analysis of *RiOLE1-LIKE* related sequences revealed that this gene is conserved in AM fungi from the Glomales and Diversisporales, but is absent from non-symbiotic Mortierellaceae and Mucoromycotina fungi, suggesting that 16:1^Δ11cis^ provides a specific function during AM colonization.

## Introduction

Mycorrhiza fungi establish symbiotic interactions with plant roots. The fungi provide mineral nutrients, particularly phosphate, to the host, in exchange for reduced carbon [1]. AM fungi establish an intricate membrane system in the cytosol of the root cortex cell, the so-called arbuscule, where nutrient and carbon exchange takes place [2]. Previously, it was believed that carbon is transported to the fungus solely in the form of carbohydrates. The fact that AM fungi lack the genes for fatty acid *de novo* synthesis, together with recent evidence that plant-derived fatty acids accumulate in the fungus, demonstrated that in addition to carbohydrates, fatty acid-containing lipids are transferred from the plant to the fungus [3–6]. In line with this scenario, expression of host specific genes (e.g. *DIS, FatM, RAM2*) involved in fatty acid and lipid synthesis is upregulated during colonization. The *DIS* gene encodes plastidial β-keotacyl-acyl carrierer (ACP) synthase I (KASI) involved in the production of palmitoyl-(16:0)-ACP [6]. *FatM* encodes a plastidial acyl-ACP thioesterase, with high activity for hydrolysis of 16:0-ACP [7,8]. After export from the plastids, fatty acids are converted into acyl-CoA thioesters. The RAM2 gene product produces sn2-acyl-glycerol form glycerol-3-phosphate and acyl-CoA, with preference for 16:0-CoA [3]. Recently, methyl-myristic acid (methyl-14:0) was shown to induce spore formation of *R. irregularis* [9]. Furthermore, myristic acid (14:0) supplementation to the medium was sufficient to support hyphal growth in axenic culture, 14:0 was taken up by the fungus and metabolized [10]. Taken together, these results suggest that a 14:0 or 16:0-containing lipid most likely is transferred from the host to the fungus where it serves as carbon source in addition to carbohydrates.

While AM fungi genomes lack the genes of fatty acid *de novo* synthesis, they do contain genes encoding fatty acid desaturases and elongases [4]. Therefore, AM fungi can desaturase and produce very long chain fatty acids from precursors such as 14:0 or 16:0 [11]. The major fatty acids in AMF are 16:0 and 16:1^Δ11cis^ (11-cis-palmitvaccenic acid; 16:1^ω5cis^). 16:1^Δ11cis^ is absent from plants. It accounts for 46 - 78 mol% of total fatty acids in spores of AMF of the Glomales including the families Acaulosporaceae, Glomaceae and Gigasporaceae, albeit some species of Glomaceae (*Glomus leptotichum, Glomus occultum*) and Gigasporacea (*Gigaspora albida, Gigaspora gigantea, Gigaspora margarita, Gigaspora rosea*) contain no or traces of 16:1^Δ11cis^. In addition to 16:1^Δ11cis^, *Rhizophagus irregularis, Glomus claroideum* and *Gigaspora roseae* spores and hyphae contain 0.2 - 3.3 mol% of 16:1^Δ9cis^ (palmitoleic acid; 16:1^ω7cis^). Glomaceae species lacking 16:1^Δ11cis^ (*Glomus leptotichum, Glomus occultum*) instead contain considerable amounts (11 - 55 %) of 16:1^Δ9cis^. Spores of *Gigaspora* spp. with low amounts of 16:1^Δ11cis^ contain high contents (38 - 48 %) of 18:1^Δ9cis^ (oleic acid) and 8 - 15 % of 20:1^Δ11cis^ (gondoic acid) [12]. Oleic acid and gondoic acid are present in low amounts in spores of *R. irregualris* and other Glomaceae [11,12]. Palmitoleic acid, oleic acid, and gondoic acid are not AM fungi-specific, because they are found in non-colonized plant roots [11,13]. AM fungi additionally contain C20, C22 and C24 fatty acids. Of these, 20:3 is the predominant one with 0.2 - 1.5 mol% in *R. irregularis* [11,13].

16:1^Δ11cis^ is only found in AM fungi but is absent from plants and from most other fungi except for low amounts in the ectomycorrhiza fungi *Pisolithus tinctorius* and *Laccaria bicolor* [14,15]. 16:1^Δ11cis^ is also found in some bacterial species, including *Cytophaga hutchinsonii* or *Lactobacillus* spp. [16,17]. In some moths, 16:1^Δ11cis^ serves as intermediate for the synthesis of sex pheromones. In *Trichoplusia ni*, 16:1^Δ11cis^ is chain-shortened to 14:1^Δ9cis^ and 12:1^Δ7cis^, the latter being reduced to the alcohol and acetylated, forming the active pheromone cis-7-dodencenyl acetate [18]. In the small ermine moths *Yponomeuta evonymella* and *Y. padella*, 11-cis-hexadecenol-acetate is directly used as sex pheromone [19].

In AM fungi, 16:1^Δ11cis^ accumulates in different glycerolipids, mostly in the storage lipid triacylglycerol, but it is also found in phospholipids [4,13]. Because of the rather unique presence in AM fungi, 16:1^Δ11cis^ and of other specific fatty acids (e.g. 20:3), these fatty acids were used as lipid biomarkers for AM fungi under laboratory and field conditions [12,20,21]. In line with this scenario, the rate of colonization with *R. irregularis* correlates with the accumulation of 16:1^Δ11cis^ in colonized roots [22–24].

The gene involved in 16:1^Δ11cis^ production in AM fungi remained unknown. Evaluation of the *R. irregularis* genome and transcriptome revealed the presence of one sequence (*RiOLE1*) with high similarity to the *Saccharomyces cerevisiae OLE1* (*oleic acid dependent1*) gene, and a second sequence (*RiOLE1-LIKE*) with lower similarity [4]. In addition, the *R. irregularis* genome contains five sequences with high similarity to the Δ12 desaturase, Δ5 desaturase and Δ6 desaturase sequences from *Mortierella alpina* [4]. To identify the gene involved in 16:1^Δ11cis^ production in *R. irregularis*, we focused on the characterization of *RiOLE1* and *RiOLE1-LIKE*. Expression in the *S. cerevisiae Δole1* mutant together with fatty acid analysis and fatty acid feeding experiments revealed that *Ri*OLE1 predominantly produces oleic acid (18:1^Δ9cis^), while *RiOLE1-LIKE* is crucial for the synthesis of 16:1^Δ11cis^. *Ri*OLE1-LIKE was subsequently introduced into transgenic tobacco and Camelina, to evaluate its capacity to produce 16:1^Δ11cis^ in a plant system, and the phylogenetic distribution of *Ri*OLE1-LIKE in symbiotic and non-symbiotic fungi was investigated.

## Experimental

### Phylogenetic analysis

For retrieving the desaturase sequences from *R. irregularis*, the four characterized *Mortierella alpina* desaturase sequences were retrieved from UniProtKB (uniprot.org) and used in BLASTp searches at the EnsemblFungi database (fungi.ensembl.org). For phylogenetic analysis of desaturases in symbiotic and non-symbiotic Mucoromycota, *RiOLE1* and *RiOLE1-LIKE* sequences were used in BLASTn searches to identify orthologs in *Gigasprora rosea, Rhizophagus cerebriforme* and *Rhizophagus diaphanus* genomes available at MycoCosm [25]. *Rhizophagus clarus, Mortierella circinelloides, Rhizophagus microsporus, Mortierella elongata* and *Lobosporangium transversale* genomes at EnsemblFungi were surveyed for *RiOLE1* and *RiOLE1-LIKE* orthologs using BLASTp. The sequences for phylogenetic analyses were aligned using MUSCLE implemented in MEGA 7, and the alignments used to create a maximum-likelihood tree with the WAG +F nucleotide substitution model. Gamma distributed rates with invariant sites (G+I) and 1000 bootstrap iterations were performed [26].

### Expression of *RiOLE1* and *RiOLE1-like* in yeast

The cDNAs of *RiOLE1* and *RiOLE1-LIKE* were amplified from *R. irregularis* RNA isolated from colonized *Lotus japonicus* roots by RT-PCR (for oligonucleotides, see Supplementary Table S1). The cDNAs were ligated into the expression vector pDR196 using the restriction enzymes *Sma*I and *Xho*I [27]. The constructs were transferred into *Saccharomyces cerevisiae* BY4741 (wild type, WT) and the *Δole1* mutant (accession Y40963) obtained from EUROSCARF, and transformed cells selected on minimal medium lacking uracil. The growth of *Δole1* was supported by addition of 1 mM oleic acid.

For complementation experiments, single colonies of *Δole1* cells containing different constructs were used to inoculate 20 mL cultures of minimal medium with or without 1 mM oleic acid/1% (v/v) Triton-X 100, and cells were grown at 33°C. Alternatively, the cells were spotted in a serial dilution on solid medium and grown at 28°C for five days.

Fatty acid feeding experiments were performed with whole *S. cerevisiae* cells. To this end, minimal medium lacking uracil was inoculated with single colonies from plates. Cells were supplemented with 1% (v/v) Triton X-100 and 1 mM of 15:0 or ^13^C_4_-16:0 fatty acid, and grown for 2 days at 28°C.

### Expression of *RiOLE1* and *RiOLE1-like* in plants

The cDNAs for *RiOLE1* and *RiOLE1-LIKE* were amplified by PCR for expression in *Nicotiana benthamiana* (for oligonucleotides, see Supplementary Table S1). *RiOLE1* was cloned into the vector p917RFPUBQExpr (Nicole Gaude, MPI Potsdam-Golm) using *Bam*HI, *Hin*dIII, and *RiOLE1-LIKE* was ligated into the vector pBin35S-DsRed using *Mlu*I, *Xho*I [28]. The constructs were transferred into *Agrobacterium tumefaciens* GV3101-pMP90 and used for transient transformation of *N. benthamiana* leaves [29]. Plants with infiltrated leaves were grown at 25°C, 60% humidity and light intensity of 250 mmol s^−1^ m^−2^ for 4 to 11 days and expression of the *DsRed* marker was evaluated with a fluorescent lamp (NightSea, Bedford, USA).

For expression in *Camelina sativa, RiOLE1* and *RiOLE1-LIKE* cDNAs were amplified by PCR (for oligonucleotides, see Supplementary Table S1) and cloned into pBinGlyBar1 (Edgar Cahoon, University of Nebraska-Lincoln) using *Eco*RI, *Xho*I and transferred into *A. tumefaciens* GV3101. *Camelina sativa* accession CAM139 (ACCID 243618; IPK Gatersleben, Germany) [30], and a high-palmitate line expressing *CpuFatB1* [31] were transformed via vacuum infiltration [32]. T1 plants were germinated and screened by spraying the plants at 5 and 12 days post germination with 0.3 % (v/v) BASTA (Bayer CropScience) containing 500 *µ*L L^−1^ Silwet L-77. Seeds from resistant plants were harvested for single-seed fatty acid analysis.

### Lipid extraction and fatty acid analysis

Lipids were extracted from yeast cultures and *N. benthamiana* leaves. Yeast cultures were centrifuged at 4000 g for 10 min and the pellet resuspended in 4 mL water and boiled for 5 min. Lipids were extracted with three volumes of chloroform/methanol (2:1, v/v) after centrifugation at 5 min for 2000 g and re-extracted two times with two volumes of chloroform each. Transformed *N. benthamiana* leaf areas were dissected and frozen in liquid nitrogen. Leaves were homogenized in a mortar under liquid nitrogen and the leaf powder was used for lipid extraction with chloroform/methanol/formic acid (1:1:0.1, v/v) and 1 M KCl/0.2 M H_3_PO_4_. The solvent of the lipid extracts were evaporated under N_2_ gas and the lipids separated by solid-phase extraction or directly used for synthesis of fatty acid methyl esters.

For solid phase extraction, lipids were dissolved in 1 mL chloroform and loaded onto silica columns (1 mL bed volume; Phenomenex, Torrance, CA, USA). Non polar lipids were eluted with chloroform, galactolipids with acetone/isopropanol 1:1 (v/v) and phospholipids with methanol. The solvents were evaporated with N_2_ gas.

Fatty acid methyl esters were synthesized from dried total lipid extracts, solid phase extraction-fractions, or single seeds of Camelina by incubation with 1 N methanolic HCl at 80°C for 30 min [33]. Methyl esters were extracted with n-hexane and 0.9% (w/v) NaCl. Camelina seeds were homogenized prior to incubation with methanolic HCl. Fatty acid methyl esters from *R. irreglaris*-colonized *L. japonicus* roots or *R. irregularis* extraradical mycelium (ERM) were prepared as previously described [7]. Fatty acid methyl esters were analyzed by gas chromatography with flame ionization detector (Agilent 7890A Plus GC; Agilent, Santa Clara, CA, USA) [4] or by gas chromatography-mass spectrometry (6975C inertXL MSD with Triple-Axis Detector with 7890A GC, Agilent) with an HP 5-MS column (Agilent J&W, 30 m, 0.25 mm diameter, 0.25 *µ*m film). Pentadecanoic acid (15:0) was used as internal standard.

The double bond positions were investigated after conversion of monounsaturated fatty acid methyl esters with dimethyldisulfide (DMDS) into bis(methylthio)-derivatives [34]. Dimethyldisulfide derivatives were analyzed by GC-MS.

### Measurement of acyl-CoAs

Acyl-CoAs were extracted from homogenized leaf material of *N. benthamiana* (20 mg) as described [35,36]. For measurement in yeast, the cell pellet of a 2 mL culture was resuspended in 200 *µ*l of the extraction buffer (1 mL isopropanol, 1 mL 50 mM KH_2_PO_4_, pH 7.2, 25 *µ*L acetic acid, 40 *µ*L 50 mg mL^−1^ fatty acid free bovine serum albumine) and homogenized in the Precellys homogenizer with glass beads. Extracted acyl-CoAs were dissolved in solvent B (H_2_O/acetonitrile; 90:10; v/v; containing 15 mM NH_4_OH) and separated on an RP8 column (Knauer Eurospher II, 150 mm, 3 mm) at a flow rate of 0.5 mL min^−1^ with the following gradient: 0 min, 100% solvent B; 5 min, 25% solvent A (acetonitrile containing 15 mM NH_4_OH)/75% solvent B; 11-13 min, 100 % solvent A; 15-18 min, 100% solvent C (H_2_O/acetontrile/formic acid, 30:70:0.1, v/v); 20-30 min, 100% solvent B. Eluted acyl-CoAs were quantified by MS/MS with electrospray ionization on an Agilent 6530 Q-TOF mass spectrometer. Acyl-CoAs were ionized in the positive mode and quantified using the neutral loss of 506.9960 (characteristic for adenosine-3’-phosphate-5’-diphosphate) using 17:0-CoA as internal standard.

## Results

### *R. irregularis* contains two genes with sequence similarity to the *S. cerevisiae* acyl-CoA desaturase OLE1

The genome of *R. irregularis* harbors seven open reading frames with sequence similarity to desaturases from *Mortierella alpina* and *Saccharomyces cerevisiae*. These genes can be divided into three clades based on amino acid sequence relationship (Figure Supplementary S1). Clade I contains sequences from *M. alpina* with specificity for Δ5 and Δ6 desaturation [37] and three *R. irregularis* genes. Clade II contains one *R. irregularis* sequence and two *M. alpina* desaturases specific for Δ12 and ω3. The *M. alpina* and *S. cerevisiae* OLE1 Δ9 desaturases belong to Clade III and show highest similarity with *R. irregularis* RIR 1928200 (designated *RiOLE1*), and the second most similar sequence is RIR 2584800 (*Ri*OLE1-LIKE).

Expression of the two genes was recorded in *Daucus carota* roots growing in tissue culture which were infected with *R. irregularis* (+ myc, intraradical mycelium, IRM) and extraradical mycelium (ERM). Semiquantitative RT-PCR with total RNA revealed that expression of *RiOLE1* and *RiOLE1-LIKE* is similar in IRM and ERM, even though *RiOLE1-LIKE* shows stronger overall expression (Supplementary Figure S2A). Therefore, expression of *RiOLE1* or *RiOLE1-LIKE* is not upregulated in the fungus during AM colonization, but the two genes are constitutively expressed.

To study the biochemical function of *R. irregularis Ri*OLE1 and *Ri*OLE1-LIKE, the two cDNAs were transferred into the fatty acid-auxotroph yeast Δ*ole1* mutant. This yeast mutant is deficient in the 18:0-CoA Δ9 desaturase and therefore can only grow after supplementation of unsaturated fatty acids such as oleic acid (18:1^Δ9cis^) to the medium. Introduction of *Ri*OLE1 or *Ri*OLE1-LIKE both resulted in complementation of the growth deficiency of Δ*ole1* (Figure 1A, B). The yeast Δ*ole1* mutant was used to examine the fatty acid desaturation products after heterologous expression of *Ri*OLE1 and *Ri*OLE1-LIKE by GC (Figure 1C). Expression of *RiOLE1-LIKE* led to the accumulation of the mycorrhiza signature fatty acids, 11-cis-palmitvaccenic acid (16:1^Δ11cis^) and vaccenic acid (18:1^Δ11cis^) (Figure 1C, Supplementary Figure S2A). On the other hand, expression of *RiOLE1* resulted in a strong increase in oleic acid (18:1^Δ9cis^) content. Expression of the two desaturases in Δ*ole1* led to a slight decrease in palmitoleic acid (16:1^Δ9cis^), and *RiOLE1* expression slightly affected the myristic acid (14:0) content. However, no desaturation product of 14:0 (14:1) was detected in any yeast strain.

**Figure 1.**
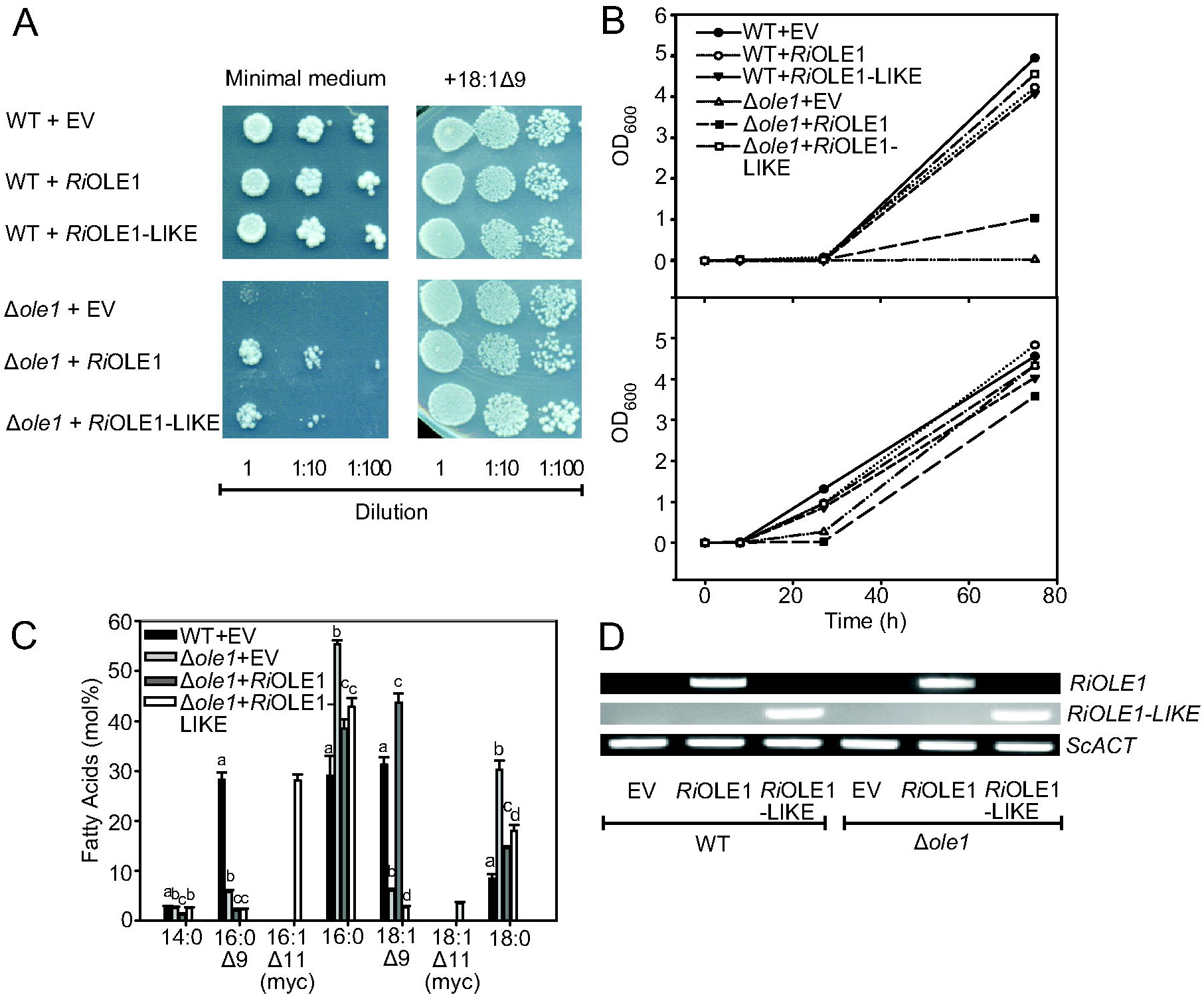
Complementation of yeast Δ*ole1* mutant growth by heterologous expression of *Rhizophagus* desaturases. *RiOLE1* or *RiOLE1-LIKE* cDNAs were expressed in *S. cerevisiae* WT and the Δ*ole1* mutant cells. Cells were grown in minimal medium with or without supplementation of oleic acid (18:1^Δ9cis^). (**A**) Spotting growth assay of yeast cultures. Cells of 10 fold dilutions were pipetted onto solid medium and grown at 28°C for five days. (**B**) Liquid cultures were inoculated from single colonies and grown at 33°C for up to three days, and the OD_600_ was recorded. (**C**) Fatty acid composition of yeast WT and Δ*ole1* cells expressing *RiOLE1* or *RiOLE1-LIKE*. Total lipids were extracted from the harvested cells and used for synthesis of fatty acid methyl ester which were quantified by GC. Bars show means ± SD. Letters indicate significant differences between treatments (ANOVA; posthoc Tukey; p ≤ 0.05; n=3). (**D**) Expression of *Ri*OLE1 or *Ri*OLE1-LIKE in yeast WT and Δ*ole1*. RNA was isolated from WT or Δ*ole1* cells transformed with empty vector (EV), *Ri*OLE1 or *Ri*OLE1-LIKE constructs. Transcript abundance was measured by RT-PCR with primers for *RiOLE1, RiOLE1-LIKE* or *ACTIN.* PCR products were separated on agarose gels and stained with ethidium bromide.

*RiOLE1* and *RiOLE1-LIKE* gene expression in the different yeast cultures was determined by semiquantitative RT-PCR (Figure 1D). *RiOLE1* and *RiOLE1-LIKE* were expressed to the same extents in the recombinant yeast cultures, indicating that the differences in fatty acid desaturation are due to different activities or substrate preferences of the enzymes.

### Determination of double bond position of mycorrhiza-signature fatty acids

The double bond position of unsaturated fatty acids can be determined after covalent modification of double bonds with dimethyldisulfide followed by GC-MS analysis, giving rise to peaks with specific retention times and fragmentation patterns. Therefore, total lipids isolated from the different yeast Δ*ole1* strains were converted into methyl esters and derivatized with dimethyldisulfide (Figure 2). Four monunsaturated fatty acids were present in the total fatty acid extracts (Supplementary Figure S3). Their fragmentation patterns allowed the identification of the double bonds at Δ9 of palmitoleic acid (16:1^Δ9is^) and oleic acid (18:1^Δ9cis^) (Figure 2A, 2C) and Δ11 for palmitvaccenic acid (16:1^Δ11cis^) and vaccenic acid (18:1^Δ11cis^) (Figure 2B, 2D). While palmitoleic acid and oleic acid accumulated in *RiOLE1* expressing cells, palmitvaccenic acid and vaccenic acid were abundant in *RiOLE1-LIKE* cells (Supplementary Figure S3).

**Figure 2.**
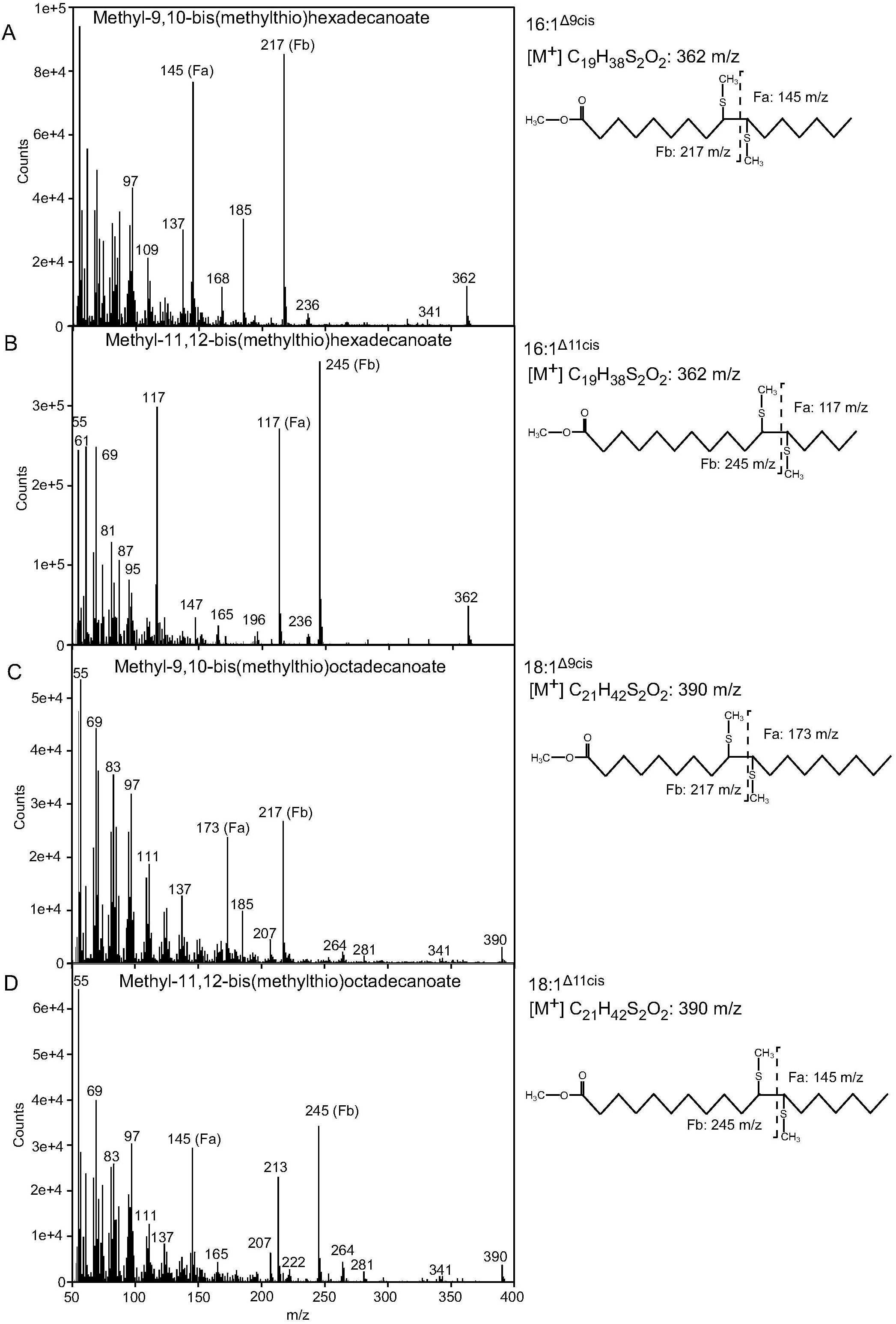
Determination of double bond position monounsaturated fatty acids isolated from *RiOLE1* or *RiOLE1-LIKE* expressing yeast. Lipids were isolated from yeast Δ*ole1* mutant cells expressing *RiOLE1-LIKE* and fatty acid converted into methyl esters. Double bonds were derivatized with dimethylsidulfide and the products analyzed by GC-MS (for chromatograms, see Supplementary Figure S2). Mass spectra are and structures are shown for **A**, 16:1^Δ9cis^; **B**, 16:1^Δ11cis^; **C**, 18:1^Δ9cis^; **D**, 18:1^Δ11cis^. F_a_, fragment a; F_b_, fragment b.

### Heterologous expression of *Ri*OLE1 and *Ri*OLE1-LIKE in transgenic plants

To characterize the two *R. irregularis* desaturases in a non-fungal background, *Agrobacterium* harboring vectors with the *Ri*OLE1 and *Ri*OLE1-LIKE cDNAs were infiltrated into *Nicotiana benthamiana* leaves for transient expression (Supplementary Figure S4). Expression of *RiOLE1* resulted in an increase in oleic acid in non polar lipids, galactolopids and phospholipids, accompanied by increased 18:2 content in non polar lipids and phospholipids (Supplementary Figure S4A). *RiOLE1-LIKE* expression in *N. benthamiana* leaves led to the accumulation of 16:1^Δ11cis^, particularly in neutral lipids and phospholipids but not galactolipids (Supplementary Figure S4B). These results are in agreement with the data obtained from heterologous expression in yeast and demonstrate that *Ri*OLE1 and *Ri*OLE1-LIKE encode a Δ9 and Δ11 desaturase, respectively.

It is known that unusual fatty acids such as 16:1^Δ11cis^ are excluded from plant membrane lipids, but they can accumulate in the seed oils of many plants [38]. We therefore expressed *RiOLE1-LIKE* using a seed-specific promoter in Camelina to study the fatty acid composition of the oil in mature seeds. Fatty acid measurement showed that 16:1^Δ11cis^ accumulated to 0.03 - 0.08 mol% in single transgenic *RiOLE1-LIKE* expressing Camelina seeds (Supplementary Table S2), while it was undetectable in WT seeds. In addition, the content of 20:1^Δ11^ increased from 12.7 mol% to 17.5 - 20.5 mol% when *RiOLE1-LIKE* was expressed. The very low amount of 16:1^Δ11cis^ in transgenic Camelina seeds might be due to the low availability of the substrate, 16:0, because Camelina seeds mostly accumulate C18 fatty acids. We therefore introduced the *RiOLE1-LIKE* construct into a high 16:0-containing Camelina line [31]. This line carries an overexpression construct for the acyl-ACP thioesterase CpuFatB1 from *Cuphea pulcherrima* resulting in the termination of fatty acid synthesis at C16, and in the accumulation of 16:0 in the seed oil from ∼11 % to ∼42 % (Supplementary Table S2). Expression of *RiOLE1-LIKE* in transgenic CpuFatB1 seeds resulted in the increase in palmitvaccenic acid content to 0.24 - 0.82 mol% (Supplementary Table S2). Likewise, the content of 20:1^Δ11^ was slightly increased, albeit total amounts were lower compared with CAM139 transformed seeds. Taken together, *RiOLE1-LIKE* expression in the seeds of Camelina resulted in the accumulation of 16:1^Δ11cis^, albeit with low amounts, and the amount was increased but still < 1 mol% in a high pamitic acid Camelina line.

### Addition of exogenous fatty acids to yeast

To examine fatty acid substrate specificity of *Ri*OLE1 and *Ri*OLE1-LIKE in more detail, fatty acids were added to growing yeast cultures. We first added a stable isotope labeled 16:0 fatty acid (^13^C_4_-16:0) to follow the incorporation of label into the different desaturation and elongation products. When ^13^C_4_-16:0 was supplied to the yeast cells, the ^13^C_4_-label accumulated in all fatty acids in WT, but was strongly reduced in 16:1^Δ9cis^ and 18:1^Δ9cis^ in the Δ*ole1* mutant where 18:0 accumulated instead (Figure 3A). Upon expression of *RiOLE1* and *RiOLE1-LIKE* in Δ*ole1*, ^13^C_4_-label accumulation in 16:1^Δ9cis^ was even further decreased. In addition, oleic acid became abundantly labeled when *RiOLE1* was expressed in Δ*ole1*, while expression of *RiOLE1-LIKE* in Δ*ole1* led to the most abundant accumulation of ^13^C_4_-label in 16:1^Δ11cis^ (54.44 mol%) and none in 18:1^Δ9cis^.

**Figure 3.**
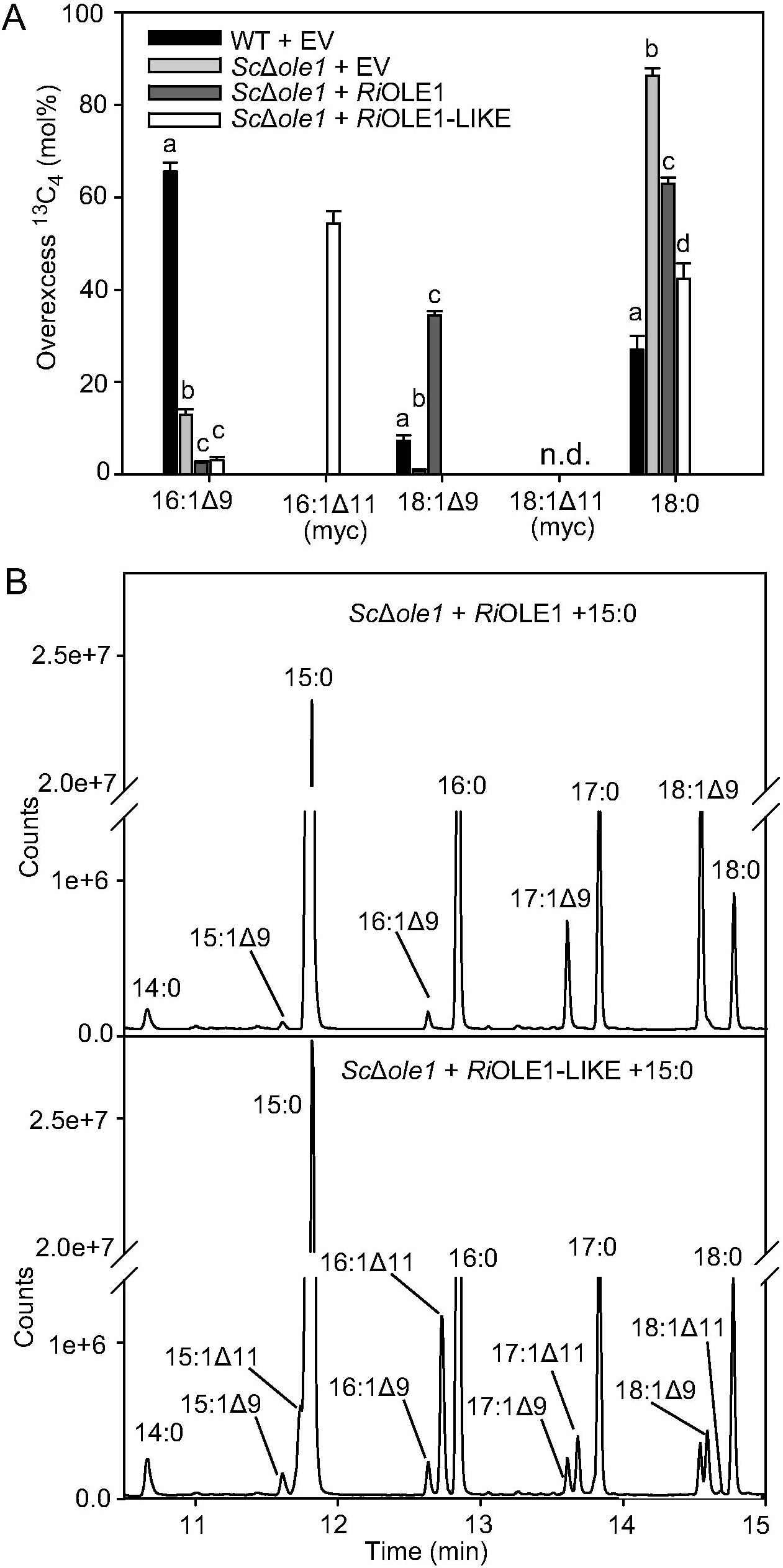
Desaturation of exogenously supplied fatty acids in yeast cells expressing *Rhizophagus* desaturases. *cerevisae* cells (WT, Δ*ole1*) expressing *RiOLE1* or *RiOLE1-LIKE* were grown for three days at 28°C in minimal medium supplemented with (**A**) 1 mM ^13^C_4_-16:0 or (**B**) 1 mM 15:0. Lipids were extracted from cell pellets and used for fatty acid analysis of methyl esters by GC-MS. (**A**) The amount of overexcess ^13^C_4_ accumulation (mol%) was quantified in the different fatty acid methyl esters. Letters indicate significant differences among treatments (ANOVA; posthoc Tukey, p ≤ 0.05, n=3). Bars represent means ± SD. n.d., not detected; EV, empty vector. (**B**) GC Chromatograms showing methyl esters of yeast lipids after 15:0 feeding.

Next, we added the odd-chain fatty acid pentadecanoic acid (15:0) to Δ*ole1* cells expressing *RiOLE1* or *RiOLE-LIKE.* 15:0 was taken up by the cells and elongated to margaric acid (17:0), and both 15:0 and 17:0 were desaturated to 15:1^9cis^ and 17:1^9cis^, respectively, in Δ*ole1* cells expressing *RiOLE1* (Figure 3B). When *RiOLE1-LIKE* was expressed in Δ*ole1*, 15:0 and 17:0 were converted into two new peaks which were tentatively identified as 15:1^Δ11cis^ and 17:1^Δ11cis^ based on their retention time.

### Acyl-CoA composition in yeast cells and plants expressing *R. irregularis* desaturases

Acyl-CoAs are important intermediates of fatty acid metabolism, and they are the substrate for the OLE1 desaturase from *S. cerevisiae*. Acyl-CoAs were quantified in yeast cells and *N. benthamiana* leaves expressing *RiOLE1* or *RiOLE1-LIKE* (Figure 4) by LC-MS/MS. In this method, acyl-CoAs containing different numbers of carbon atoms or double bonds in the acyl chain can be separated and quantified using an internal standard. However, it is not possible to separate isomeric monounsaturated acyl-CoAs containing the doubIe bond at different positions. In yeast Δ*ole1*, 16:1-CoA and 18:1-CoA decreased while 16:0-CoA, 18:0-CoA and 20:0-CoA accumulated, compared to WT (Figure 4A). Expression of *RiOLE1-LIKE* in Δ*ole1* led to a very strong increase in 16:1-CoA (43.6 mol%) and to a lesser increase in 18:1-CoA (7.6 mol%) while the amount of 20:0-CoA (2.9 mol%) was decreased. On the other hand, 18:1-CoA increased most predominantly when *RiOLE1* was expressed (26.7 mol%).

Expression of *RiOLE1-LIKE* in *N. benthamiana* leaves led to the accumulation of 16:1-CoA (16.8 mol%) accompanied with a decrease in 16:0-CoA (41.1 mol%) that was not detected in control leaves (Figure 4B). Expression of *RiOLE1* led to the increase in 18:1-CoA to 5.2 mol%, which remained low in the control (1.3 mol%) and *RiOLE1-LIKE* (2.2 mol%). Taken together, these results are in agreement with a substrate/product relationship of 18:0-CoA/18:1-CoA and 16:0-CoA/16:1-CoA for *Ri*OLE1 and *Ri*OLE1-LIKE, respectively, suggesting that *Ri*OLE1 and *Ri*OLE-LIKE are acyl-CoA desaturases.

**Figure 4.**
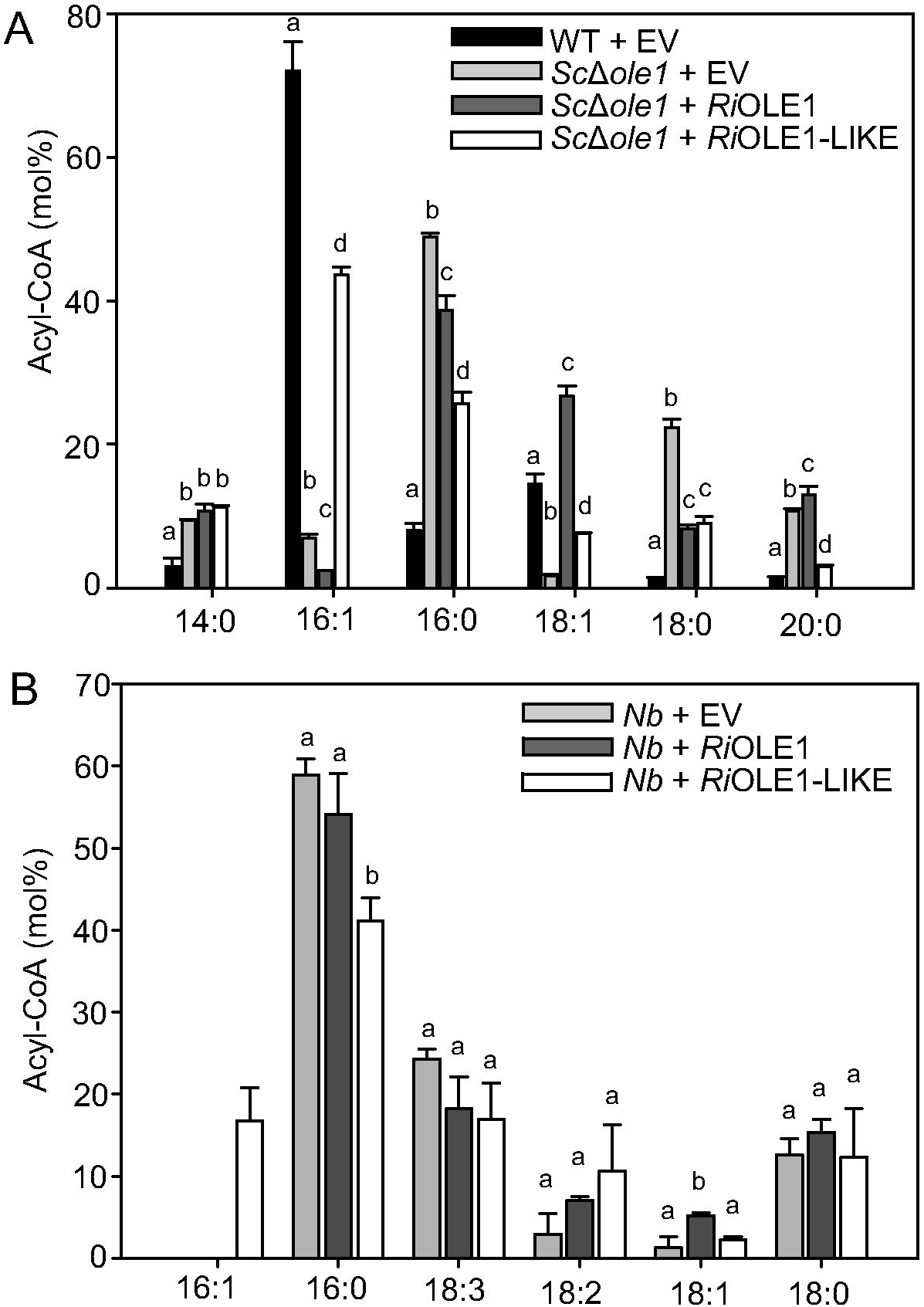
Acyl-CoA composition in transformed yeast cells and *N. benthamiana* leaves expressing *Rhiozphagus* desaturases. Acyl-CoAs were measured by LC-MS/MS. (**A**) Acyl-CoAs in yeast cells (WT, Δ*ole1*) expressing empty vector (EV) control, *RiOLE1* or *RiOLE1-LIKE*. (**B**) Acyl-CoAs in *N. benthamiana* leaves transiently expressing EV control, *RiOLE1* or *RiOLE1-LIKE*. Letters indicate significant differences among treatments (ANOVA; posthoc Tukey; p ≤ 0.05. Bars represent means ± SD (n = 3). *Sc, Saccharomyces cerevisiae*; *Nb, Nicothiana benthamiana*; n.d., not detected.

### The *OLE1-LIKE* genes are conserved in AM Glomeromycotina, but not in non-symbiotic Mucoromycota

AM fungi of the Glomeromycotina likely evolved from saprotrophic Mucoromycota [39]. To study the evolution of *Ri*OLE1 and *Ri*OLE1-LIKE sequences and their distribution in the different fungal phyla, the two desaturase sequences from *R. irregularis* were used to identify orthologs in other Glomeromycotina as well as non-mycorrhizal Mucoromycota using protein Blast searches. Protein sequences of several fungal species were retrieved and used to assemble a phylogenetic tree (Figure 5). Two major clades were identified, one clade that compromises sequences similar to *Ri*OLE1-LIKE that were present in all five representatives of the Glomeromycotina. These sequences were more similar to each other than to the sequences organized in the second clade representing the sequences highly similar to *Ri*OLE1. Sequences similar to *Ri*OLE1 were found in all six Glomeromycotina species and in the non-symbiotic Mortierellaceae (*M. alpina, M. elongata, L. transversale*) as well as in the non-symbiotic Mucoromycotina (*M. circinelloides, R. microsporus*). Thus, OLE1-LIKE is conserved in symbiotic Glomales and Diversisporales, but not in non-symbiotic fungi, suggesting that it evolved to fulfill a specific function in AM fungi.

**Figure 5.**
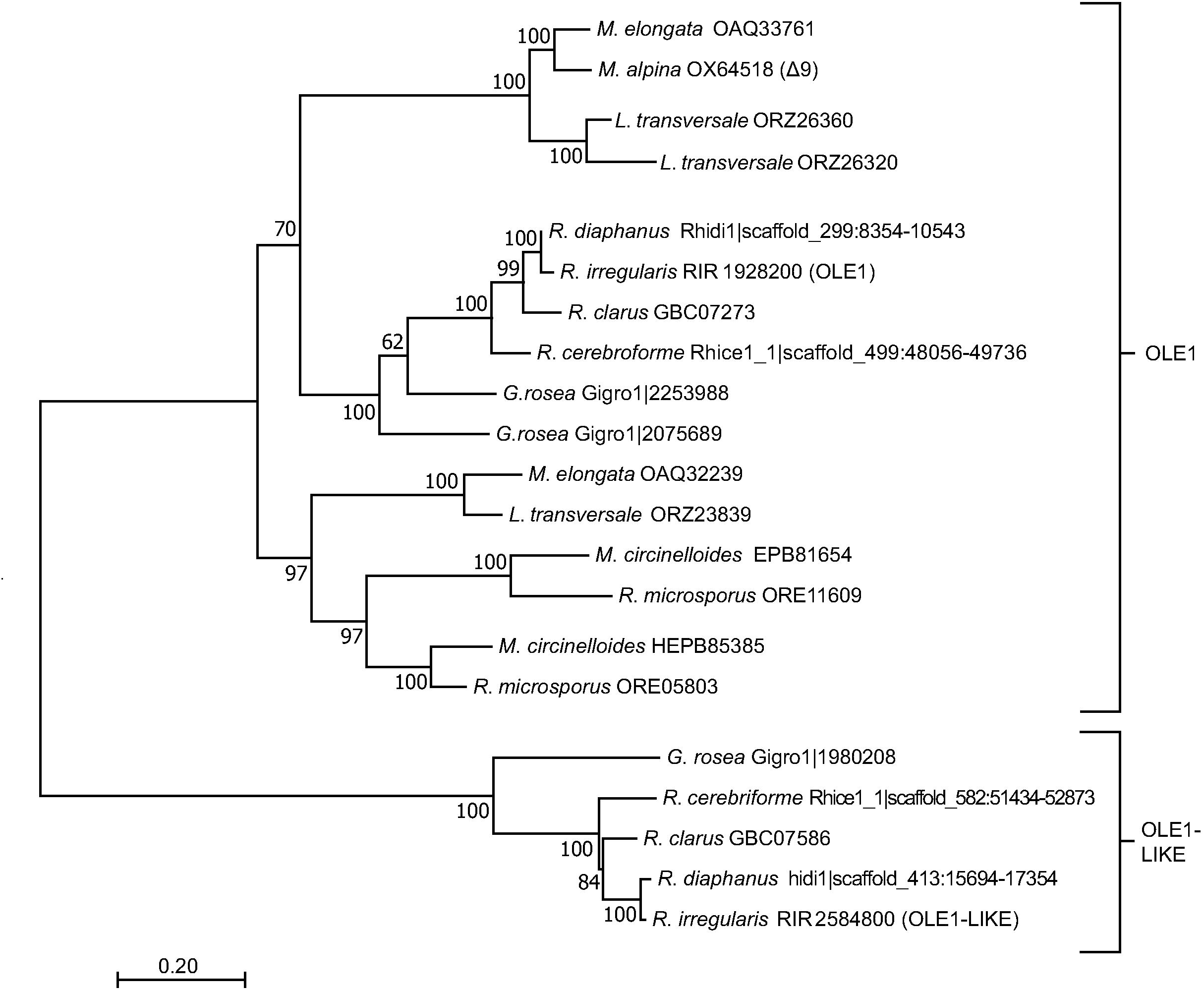
Phylogenetic tree of OLE1 and OLE1-LIKE sequences from Mucoromycota. OLE1 and OLE1-LIKE from *R. irregularis* were used to identify orthologs in species of other Glomeromycotina, Mortierellamycotina and Mucoromycotina [25]. Amino acid sequences were aligned using MUSCLE with MEGA 7 and a maximum-likelihood tree constructed with 1000 bootstrap iterations as depicted on the nodes.

## Discussion

### *Ri*OLE1 and *Ri*OLE1-LIKE are front-end desaturases

Desaturases are classified based on their ability to either recognize the methyl (ω) or carboxyl (Δ) end of fatty acids for insertion of the double bond [40]. The double bond position of fatty acids can be identified after derivatization with dimethyldisulfide. Thus, four monounsaturated fatty acids were detected in Δ*ole1* cells expressing *RiOLE1-LIKE*, i.e. 16:1^Δ9cis^, 16:1^Δ11cis^, 18:1^Δ9cis^, 18:1^Δ11cis^ (Figure 2, Supplementary Figure S3). The fatty acids 16:1^Δ9cis^ and 18:1^Δ9cis^ are also present in the Δ*ole1* control and are therefore derived from residual activity of *Sc*OLE1 rather than from *Ri*OLE1-LIKE. 16:1^Δ11cis^ could in principle be derived from front end (Δ11) or methyl end (ω5) desaturation. However, 18:1^Δ11cis^ must be derived from Δ11 desaturation. This fatty acid cannot be the product of 16:1^Δ11cis^ elongation by addition of two carbons at the carboxyl end, which would have resulted in 18:1^Δ13cis^ production. Similarly, ω5 desaturation of 18:0 by *RiOLE1-LIKE* would also have resulted in the production of 18:1^Δ13cis^. Likewise, upon *RiOLE1* expression in Δ*ole1*, only 16:1^Δ9cis^ and 18:1^Δ9cis^ were produced. While 18:1^Δ9cis^ could in principle be produced via Δ9 or ω9 desaturation, 16:1^Δ9cis^ is clearly derived from Δ9 desaturation of 16:0, because ω9 desaturation would result in 16:1^Δ7cis^ production, and elongation of a putative 14:1^Δ7cis^ precursor is not likely as it was not detected.

These conclusions are corroborated by the fatty acid analysis of yeast Δ*ole1* cultures after 15:0 feeding (Figure 3). In this experiment, 15:1^Δ9^ and 17:1^Δ9^ were strongly increased after expression of *RiOLE1* compared with the control. Expression of *Ri*OLE1-LIKE in Δ*ole1* cells resulted in the accumulation of 15:1^Δ11^ and 17:1^Δ11^ from 15:0. Taken together, these data demonstrate that *Ri*OLE1 and *Ri*OLE1-LIKE are front-end desaturases that count carbon atoms from the carboxyl-terminus for insertion of the double bond at positions Δ9 or Δ11, respectively.

### *Ri*OLE1 and *Ri*OLE1-LIKE likely are acyl-CoA desaturases with specifies for long chain fatty acids

*Ri*OLE1 and *Ri*OLE1-LIKE show sequence similarity with the yeast acyl-CoA desaturase OLE1 (Supplementary Figure S1) and can functionally replace OLE1 in the Δ*ole1* mutant, strongly promoting growth in the absence of exogenous unsaturated fatty acids (Figure 1). In similar approaches, desaturases from rat or *Rhodosporidium* with sequence similarity to OLE1 were characterized [41,42]. Expression of *RiOLE1* and *RiOLE1-LIKE* leads to alteration in acyl-CoA composition in yeast and plants with a clear substrate to product relationship (Figure 4). While *RiOLE1* expression resulted in an increase in 18:1-CoA and a decrease in 18:0-CoA, expression of *RiOLE1-LIKE* caused an increase in 16:1-CoA and a reduction in 16:0-CoA, both compared with Δ*ole1.* Therefore, *Ri*OLE1 and *Ri*OLE1-LIKE likely are acyl-CoA desaturases.

Expression of *RiOLE1* in plants (*N. benthamiana* leaves, Camelina seeds) led to a very low production of 16:1^Δ11cis^. (Supplementary Table S1, Supplementary Figure S4). In plants, most desaturation reactions take place on lipid-bound acyl groups, i.e phosphatidylcholine at the ER, before acyl groups are redistributed to other membrane lipids in the leaves or to triacylglycerol in the seeds. Introduction of heterologous acyl-CoA desaturases into plants oftentimes results in low production of the new unsaturated fatty acid because it has to be channeled between the phosphatidylcholine and CoA thioester pools, and this redistribution is slow [43].

Furthermore, addition of exogenous 15:0 resulted in the accumulation of the respective desaturation products (15:1^Δ11cis^, 17:1^Δ11cis^ for *RiOLE1-LIKE;* 15:1^Δ9cis^, 17:1^Δ9cis^ for *RiOLE1*; Figure 3B) indicating that the two enzymes are to a certain degree promiscuous in their substrate specificity and accept a range of long-chain (C15-C18) fatty acids.

Feeding of ^13^C_4_-labeled 16:0 revealed the accumulation of the ^13^C_4_-label in all endogenous yeast fatty acids (Figure 4A). Yeast OLE1 is able to desaturate 16:0 and 18:0 [41]. When ^13^C_4_-16:0 was offered, ^13^C_4_-label accumulated in 18:0 (27.0 mol%) in yeast WT. This was sufficient as substrate for *Sc*OLE1 to produce ^13^C_4_-18:1^Δ9cis^, even though this product only accumulated to 7.4 mol%. Using the same conditions, no ^13^C_4_-label was found in 18:1^Δ11cis^ in Δ*ole1* expressing *RiOLE1-LIKE*, even though sufficient substrate was available (42.5 mol% of ^13^C_4_-18:0). Therefore, *Ri*OLE1-LIKE has much poorer activity on 18:0 than yeast *Sc*OLE1, and consequently *Ri*OLE1-LIKE has a strong preference for the 16:0 substrate (Figure 1C, 2D). 18:0 desaturation in *R. irregularis* is probably foremost conducted by *Ri*OLE1, in accordance with the accumulation of ^13^C_4_-18:1^Δ9cis^ (34.4 mol%) while only miniature amounts of ^13^C_4_-16:1^Δ9cis^ (2.6 mol%) were detected upon *RiOLE1* expression. Functional differentiation of acyl-CoA desaturases was reported before for mice where four desaturases exist, three act on 16:0-CoA and 18:0-CoA and one is specialized for acyl-CoAs with fatty acids < 16:0 [44].

### OLE1-LIKE evolved in AM fungi

OLE1-LIKE is conserved in the Glomerales *R. clarus, R. irregularis, R. diaphanus, R. cerebriforme* and in *G. rosea* (Diversisporales) but not in the other non-symbiotic, non-mycorhizal Mucoromycota (Figure 5). The Mucoromycota species only contain OLE1 orthologs. A characteristic feature of Glomeromycotina are their large genomes with high numbers of duplicated genes, compared to Mortierellamycotina and Mucorocmycotina [25]. Therefore, the *OLE1-LIKE* gene could originate from gene duplication of *OLE1* and neofunctionalization in Glomeromycotina and might have evolved to fulfill a specific function during the obligate mutualistic AM symbiosis. The expression pattern of *RiOLE1-LIKE* in IRM and ERM (Supplementary Figure S2A) is consistent with the occurrence of 16:1^Δ11cis^ in non polar storage lipids and membrane-forming phospholipids in IRM and ERM, even though considerably less 16:1^Δ11cis^ is found in the IRM [13].

### Possible functions of 16:1^Δ11cis^ in AM fungi

Interestingly, overexpression of the *S. saccharomyces* Δ9 desaturase (OLE1) or of an insect Δ11 desaturase, which resulted in the accumulation of 16:1 ^Δ9cis^ and 16:1^Δ11cis^ in transgenic *Nicotiana tabacum* leaves, respectively, led to the accumulation of lipoxygenase products, in particular cis-3-hexenal [45]. The same effect was observed when the two fatty acids were exogenously applied to *N. tabacum* leaves. The authors concluded that the presence of unusual monounsaturated C16 fatty acids stimulated the lipoxygenase pathway thereby contributing to enhanced resistance against pathogens. In line with this finding, overexpression of *S. cerevisiae* OLE1 in tomato or in eggplant increased the resistance against powdery mildew (*Erysiphe polygoni*) and Verticillium wilt (*Verticillium dahliae*) [46,47]. Interestingly, 16:1^Δ9cis^ and to a lesser extent 18:1^Δ9cis^, inhibited growth of *Verticillium dahliae* at *µ*M concentrations. Therefore, it is possible that monounsaturated C16 fatty acids, in particular 16:1^Δ11cis^, which accumulates in membrane and storage lipids, protect AM fungal spores and hyphae in the soil against attack by bacteria or other fungi. Furthermore, the unusual 16:1^Δ11cis^ fatty acid could be involved in recognition of the AM fungi by the plant host. Membrane lipids of pathogenic fungi are recognized by the plant host as shown by the elicitation of defense responses after application of ergosterol [48,49]. 16:1^Δ11cis^ possibly fulfills an opposite function during symbiosis, because it might be perceived by the host as a symbiotic signature to avoid the host’s immune response.

On the other hand, certain fatty acid combinations of membrane lipids are elicitors in plant-pathogen interactions, as shown by 18:2, 18:1 and 16:0-containing phosphatidylcholine and phosphatidylethalonamine from female *Sogatella furcifera* eggs [50]. Acyl editing in IRM of *R. irregularis* suggests that 16:1^Δ11cis^ could also be perceived by the host as potential pathogenic elicitor as *R. irregularis* alters its phospholipid fatty acid composition and suppresses di-16:1 and 24:1 when growing intraradically [4]. Therefore, 16:1^Δ11cis^ biosynthesis could have been evolved in Glomeromycota for survival in the hostile soil environment during asymbiotic growth and it might be beneficial during host perception and symbiotic growth.

## Abbreviations

ACP: acyl carrier protein;
AM: arbuscular mycorrhiza;
CoA: coenzyme A;
ERM: extraradical mycelium;
IRM: intraradical mycelium;
WT: wild type.

Fatty acids are abbreviated as X:Y, where X depicts the number of carbon atoms, and Y the number of double bonds in the acyl chain.

## Author Contribution

M.B., E.B.C. and P.D. conceived the project. M.B. and P.D. designed the research. M.B. performed the experiments. M.B. and P.D. analyzed data and wrote the article.

