## Supplemental Material for "Palmitvaccenic acid (Δ11-cis-hexadecenoic acid) is synthesized by an OLE1-like desaturase in the arbuscular mycorrhiza fungus *Rhizophagus irregularis*"

**Supplementary data**

**Supplementary Table S1 Oligonucleotides used for cloning and RT-PCR**

| Oligonucleotide | Sequence (5'-3')<br>(restriction sites are underlined) | PCR Target | Reaction conditions |
|---|---|---|---|
| Heterologous expression in *S. cerevisiae* | | | |
| FW (bn3075)<br>RW (bn3083) | TATCCCGGGATGGTGGCACAGGCAACATT<br>ATACTCGAGTTATTTGGTTTTCCTATGTT | *RiOLE1* cDNA | 98°C-30 s, 10 cycles: (98°C-7 s, 58°C-20 s, 72°C- 32 s), 25 cycles: (98°C-7 sec, 68°C -20 sec, 72°C- 32 s)25x, 72°C-120 sec |
| FW (bn3077)<br>FW (bn3084) | TATCCCGGGATGGCTGCTGCGCCAGTAAA<br>ATACTCGAGCTATTCTTCCTTCTGACTCT | *RiOLE1-LIKE* cDNA | 98°C-30 s, 10 cycles: (98°C-7 s, 63°C-20 s, 72°C- 32 s), 25 cycles: (98°C-7 sec, 72°C -20 sec, 72°C- 32 s), 72°C-120 sec |
| Heterologous expression in *N. benthamiana* | | | |
| FW (bn3214)<br>RW (bn3076) | TATGGATCCATGGTGGCACAGGCAACATT<br>ATAAAGCTTTTATTTGGTTTTCCTATGTT | *RiOLE1* cDNA | 98°C-30 s, 10 cycles: (98°C-7 s, 58°C-20 s, 72°C- 32 s), 25 cycles: (98°C-7 sec, 68°C -20 sec, 72°C- 32 s), 72°C-120 sec |
| FW (bn3212)<br>RW (bn3213) | TATACGCGTATGGCTGCTGCGCCAGTAAA<br>ATACTCGAGCTATTCTTCCTTCTGACTCT | *RiOLE1-LIKE* cDNA | 98°C-30 s, 10 cycles: (98°C-7 s, 63°C-20 s, 72°C- 32 s), 25 cycles: (98°C-7 sec, 72°C -20 sec, 72°C- 32 s), 72°C-120 sec |
| Heterologous expression in *C. sativa* | | | |
| FW (bn3216)<br>RW (bn3211) | TATGAATTCATGGTGGCACAGGCAACATT<br>ATACTCGAGTTATTTGGTTTTCCTATGTT | *RiOLE1* cDNA | 98°C-30 s, 10 cycles: (98°C-7 s, 42°C-20 s, 72°C- 32 s), 25 cycles: (98°C-7 sec, 54°C -20 sec, 72°C- 32 s), 72°C-120 sec |
| FW (bn3217)[1]<br>RW (bn3213) | TATGGTCTCGAATTCATGGCTGCTGCGCCAGTAAA<br>ATACTCGAGCTATTCTTCCTTCTGACTCT | *RiOLE1-LIKE* cDNA | 98°C-30 s, 10 cycles: (98°C-7 s, 63°C-30 s, 72°C- 32 s), 25 cycles: (98°C-7 sec, 72°C -30 sec, 72°C- 32 s), 72°C-120 sec |
| Semiquantitative RT-PCR | | | |
| FW (bn3551)<br>RW (bn3552) | CCACCACTGCTGAAAGAGA<br>ACCTTCATGGAAGATGGAGC | *ScACT* cDNA | 95°C-120 s, 30 cycles: (94°C-30 s, 56°C -40 s, 72°C- 40 s), 72°C-10 min |
| FW (bn3151)<br>RW (bn3152) | TTATGCGGCTTTACTCCGT<br>ATGATTTACCACCAGGATGTTC | *RiOLE1* cDNA | 95°C-120 s, 30 cycles: (94°C-30 s, 56°C -40 s, 72°C- 40 s), 72°C-10 min |
| FW (bn3153)<br>RW (bn3154) | GGGCTGGCGAACAACTTTA<br>GTAAGTGATGACGGGCAATGA | *RiOLE1-LIKE* cDNA | 95°C-120 s, 30 cycles: (94°C-30 s, 56°C -40 s, 72°C- 40 s), 72°C-10 min |
| FW (bn1942)<br>RW (bn1943) | TGTCCAACCGGTTTTAAAGT<br>AAAGCACGTTTGGCGTACAT | *RiαTUBULIN* | 95°C-120 s, 30 cycles: (94°C-30 s, 56°C -40 s, 72°C- 40 s), 72°C-10 min |

[1]= For site-directed ligation of *RiOLE1-LIKE* into pBinGlyBar1 for expression in *C. sativa*, a *Bsa*I restriction site was added upstream of the *Eco*RI site and the *Eco*RI sticky-overhang created by *Bsa*I restriction.

**Supplementary Table S2 Quantification of fatty acids in transgenic Camelina seeds expressing the *R irregularis* desaturase *RiOLE1-LIKE***

The table shows fatty acid contents (in mol%) for Camelina CAM139 or CpuFatB1 backgrounds. Values are means ± SD for seeds of the non-transgenic lines CAM 139 and CpuFatB1. For the *Ri*OLE1-LIKE expressing lines, the results of single seed analysis from three independent lines is presented.

| Genotype | 14:0 | 16:1Δ9 | **16:1Δ11 (myc)** | 16:0 | 18:1Δ9 | 18:1 | 18:2 | 18:3 | 18:0 | 20:1Δ11 | 20:1 | 20:2 | 20:0 |
|---|---|---|---|---|---|---|---|---|---|---|---|---|---|
| | Fatty Acids (mol%) | | | | | | | | | | | | |
| CAM139 | 0.21± 0.04 | 0.20± 0.02 | **0.00± 0.00** | 11.33± 0.60 | 8.49± 1.03 | 1.14± 0.36 | 22.55± 0.93 | 36.43± 5.77 | 2.98± 0.43 | 12.73± 3.06 | 0.86± 0.30 | 1.51± 0.49 | 1.57± 0.75 |
| CAM139 + *Ri*OLE1-LIKE 1 | 0.25 | 0.32 | **0.03** | 16.78 | 6.94 | 1.87 | 33.42 | 11.08 | 4.70 | 17.48 | 1.13 | 4.20 | 1.80 |
| CAM139 + *Ri*OLE1-LIKE 2 | 0.24 | 0.47 | **0.05** | 17.47 | 3.19 | 2.27 | 37.44 | 6.65 | 4.91 | 20.46 | 1.93 | 2.43 | 2.47 |
| CAM139 + *Ri*OLE1-LIKE 3 | 0.32 | 0.49 | **0.08** | 17.68 | 5.96 | 3.24 | 34.59 | 7.68 | 4.36 | 19.39 | 1.85 | 2.37 | 1.98 |
| CpuFatB1 | 1.86± 0.90 | 0.15± 0.09 | **0.00± 0.00** | 42.34± 0.16 | 4.36± 0.44 | 1.06± 0.18 | 23.43± 0.39 | 16.48± 3.54 | 5.04± 1.33 | 2.21± 0.67 | 0.40± 0.01 | 0.56± 0.12 | 2.10± 0.89 |
| CpuFatB1 + *Ri*OLE1-LIKE 1 | 1.93 | 0.08 | **0.82** | 42.32 | 7.95 | 0.66 | 22.04 | 9.13 | 6.74 | 3.23 | 0.38 | 0.51 | 4.21 |
| CpuFatB1 + *Ri*OLE1-LIKE 2 | 3.35 | 0.13 | **0.57** | 41.27 | 7.95 | 0.90 | 23.31 | 7.68 | 6.67 | 3.04 | 0.51 | 0.71 | 3.92 |
| CpuFatB1 + *Ri*OLE1-LIKE 3 | 3.48 | 0.14 | **0.24** | 42.86 | 4.90 | 1.19 | 22.10 | 9.45 | 6.25 | 4.16 | 0.48 | 0.83 | 3.94 |

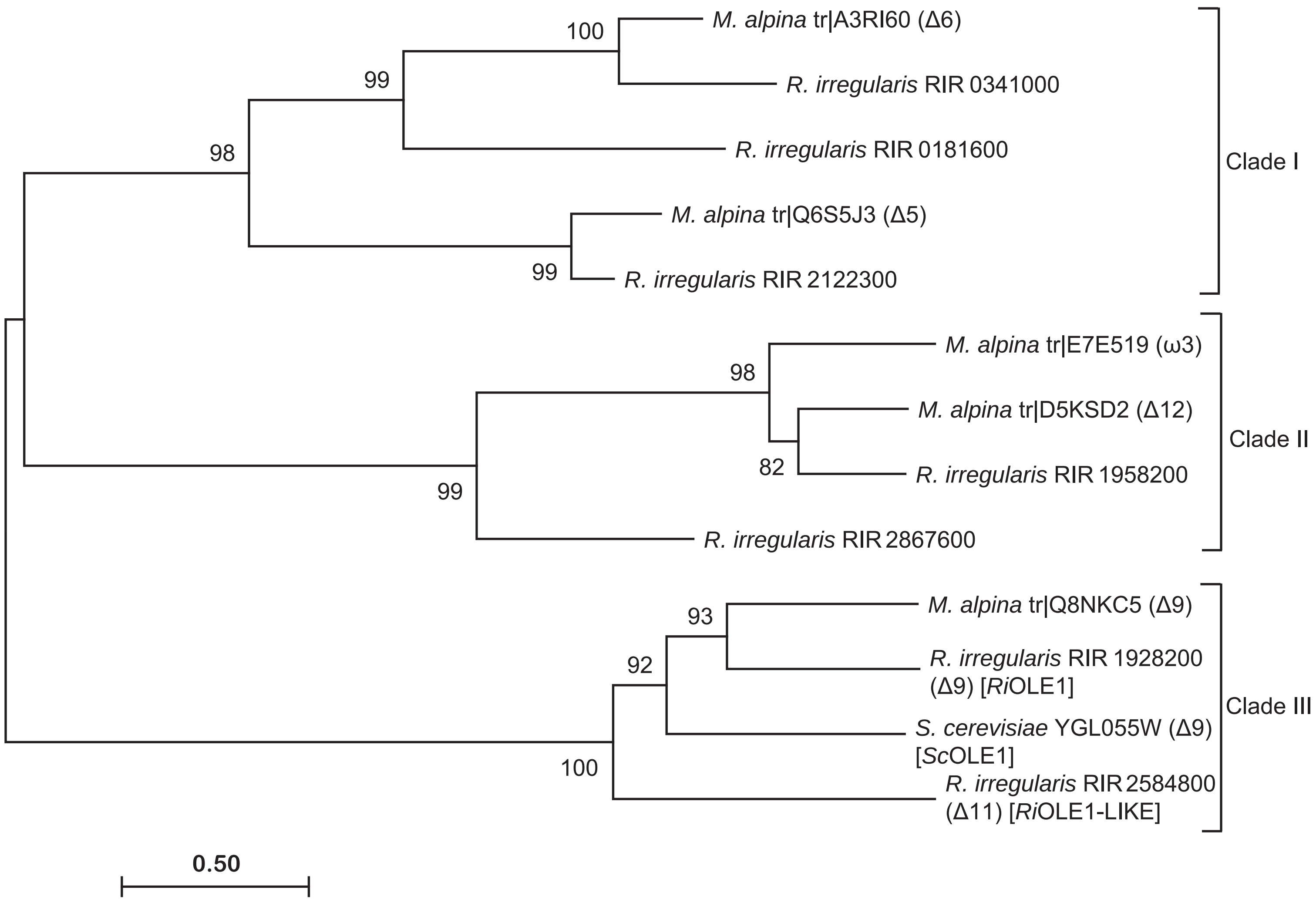


**Supplementary Figure S1. Phylogenetic tree of desaturases from *R. irregularis*, *S. cerevisae* and *M. alpina*.** Amino acids sequences were aligned using MUSCLE implemented in MEGA 7.0 and a maximum likelihood tree constructed. The branch numbers show bootstrap values (1000 iterations). The fatty acid position of desaturation by the respective desaturase is indicated in brackets. The *M. alpina* desaturase annotation was taken from [37]. *M. alpina*, *Mortierella alpina*; *R. irregularis*, *Rhizophagus irregularis*; *S. cerevisiae*, *Saccharomyces cerevisiae*.

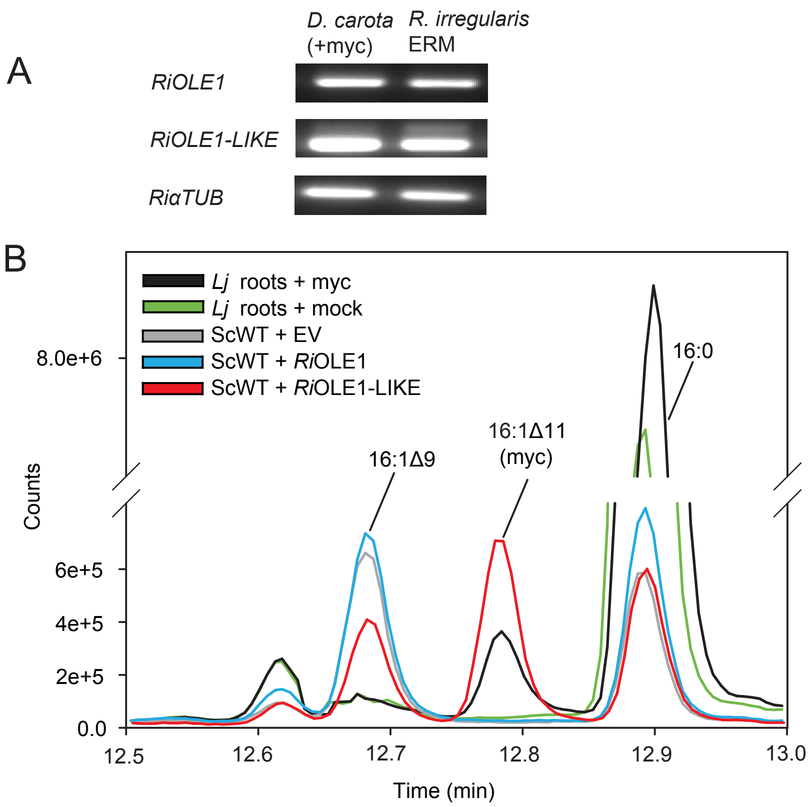


**Supplementary Figure S2. Expression of RiOLE1 and RiOLE1-LIKE and accumulation of palmitvaccenic acid (16:1Δ11cis) in plant roots colonized with *R. irregularis***
(**A**) Semiquantitative RT-PCR of *Daucus carota* roots colonized by *Rhizophagus irregularis* (+myc) and the connected extraradical mycelium (ERM). RNA from roots (+myc) and ERM was isolated from a single split-petri dish harboring an axenic, infected *D. carota* root-culture on one side and outgrowing ERM on the other side [51]. (**B**) Overlay of GC-MS chromatograms of fatty acids methyl esters from mycorrhiza-colonized and non-colonized *Lotus japonicus* roots, and yeast cells (WT) expressing RiOLE1 or RiOLE1-LIKE. The mycorrhiza-signature fatty acid 16:1Δ11cis is only present in mycorrhiza colonized roots and yeast expressing RiOLE1-LIKE. Lj, *Lotus japonicus*, Sc, *Saccharomyces cervisiae*.

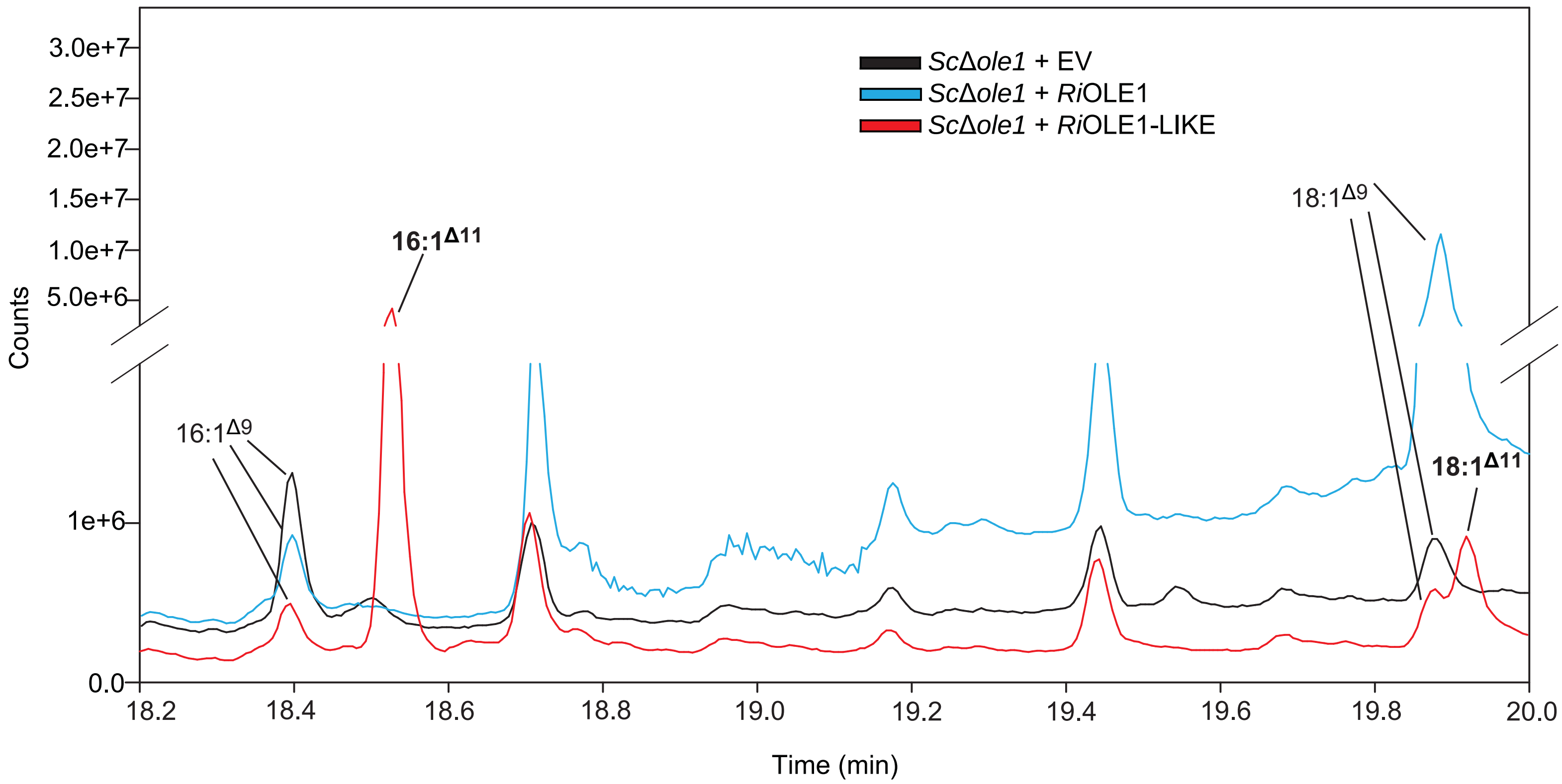


**Supplementary Figure S3. Determination of double bond positions in monounsaturated fatty acids from yeast cells expressing RiOLE1 or RiOLE1-LIKE**
Overlay of GC-MS chromatograms of dimethyldisulfide adducts of fatty acid methyl esters derived from the yeast Δole1 mutant transformed with RiOLE1 or RiOLE1-LIKE. Mass spectra and structures of the four labeled peaks are shown in Figure 3. The mycorrhiza-signature fatty acids 16:1Δ11cis and 18:1Δ11cis are highlighted in bold.

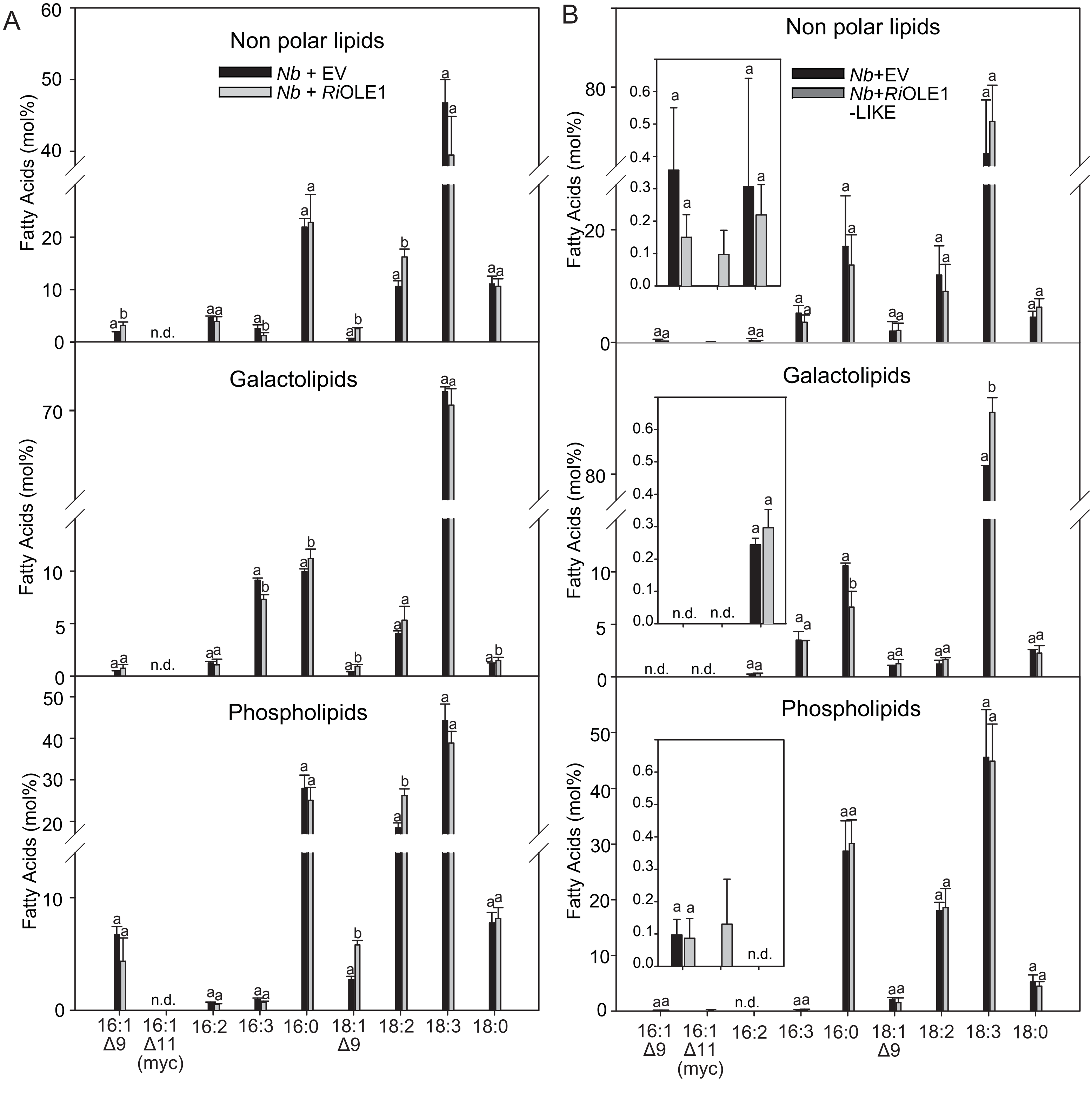


**Supplementary Figure S4. Fatty acid composition of lipid fractions from *N. benthamiana* leaves transiently expressing *Rhiozphagus* desaturases.**

Lipids isolated from *N. benthamiana* control leaves or leaves expressing RiOLE1-LIKE were separated into different fractions (non polar lipids, galactolipids, phospholipids) by solid phase extraction. Fatty acid methyl esters were produced and quantified by GC-MS. (**A**) Distribution fatty acids after expression of RiOLE1. Letters indicate significant differences among treatments (ANOVA; posthoc Tukey; $p \leq 0.05$; n=4). (**B**) Distribution fatty acids after expression of RiOLE1-LIKE. Letters indicate significant differences among treatments (ANOVA; posthoc Tukey; $p \leq 0.05$; EV, n=3 RiOLE1-LIKE, n=4). The insets show the amounts of 16:1Δ9, 16:1Δ11(myc) and 16:2 at higher scale. Bars represent means ± SD. Nb, *Nicotiana benthamiana*; n.d., not detected; EV, empty vector.
